# Prenatal murine skeletogenesis partially recovers from absent skeletal muscle as development progresses

**DOI:** 10.1101/2022.01.01.473639

**Authors:** V. Sotiriou, S. Ahmed, N. C. Nowlan

## Abstract

Skeletal muscle contractions are critical for normal growth and morphogenesis of the skeleton, but it is unclear how the detrimental effects of absent muscle on the bones and joints change over time. Joint size, shape and cavitation, and rudiment length and mineralisation were assessed in multiple rudiments at two developmental stages (Theiler Stage (TS)24 and TS27) in the splotch-delayed “muscleless limb” mouse model and littermate controls. As development progressed, the effects of absent muscle on all parameters except for cavitation become less severe. All major joints in muscleless limbs were qualitatively and quantitatively abnormal in shape at TS24, while, by TS27, most muscleless joint shapes were normal, or nearly normal. In contrast, any joints which were fused at TS24 did not cavitate by TS27. Therefore, recovery in joint shape over development occurred despite absent cavitation. Mineralisation showed the most pronounced changes between TS24 and TS27 in the muscleless limbs. At TS24, all muscleless rudiments studied had less mineralisation than the controls, while at TS27, muscleless limb rudiments had either the same or significantly *more* mineralisation than controls of the same age. We conclude that the effects of absent muscle on prenatal murine skeletogenesis are most pronounced in early skeletal development and reduce in severity prior to birth. Understanding how mammalian bones and joints continue to develop in an environment without muscle contractions, but with mechanical stimulation due to the movement of the mother, provides important insights into conditions affecting human babies such as developmental dysplasia of the hip and arthrogryposis.

## Introduction

Fetal movements play an important role in prenatal skeletal development (Nowlan, 2015). When skeletal muscle in animal models is absent, reduced in volume or non-contractile, skeletal rudiments tend to be shorter, with decreased mineralisation and malformed joints (Bridglal et al., 2021; Brunt et al., 2016; Kahn et al., 2009; Khatib et al., 2021; Nowlan et al., 2008; Nowlan et al., 2010; Nowlan et al., 2014; Pierantoni et al., 2021; Roddy et al., 2011; Sotiriou et al., 2019). Vertebral segmentation, vertebral shape and intervertebral disc formation are also dependent on muscle forces (Levillain et al., 2019; Levillain et al., 2021; Rolfe et al., 2017). Not all bones are affected equally by compromised skeletal muscles, with the bones and joints of the forelimb being more severely affected than those of the hindlimb (Kahn et al., 2009; Nowlan et al., 2010; Sotiriou et al., 2019). Unequal stimulation of the muscleless limbs from passive movements has been proposed as a possible mechanism underlying the differential effects of absent muscle on skeletal development (Nowlan et al., 2012). While it is clear that skeletal muscle contractions are critical for normal growth and morphogenesis of most bones in the limbs and spine, the effects of abnormal skeletal muscle on mammalian skeletal development over time *in utero* have not been characterised in detail.

A small number of prior studies have investigated the temporal effects of mechanical forces due to skeletal muscle contractions over development. Very early stages of skeletal development occur independently of skeletal muscle contractions, including interzone formation (Kahn et al., 2009; Mikic et al., 2000), joint morphogenesis prior to cavitation (Nowlan and Sharpe, 2014) and notochord involution (Levillain et al., 2021). The authors recently showed that the time window between embryonic days four and seven is the most important period for muscle forces in terms of the effects on hip joint development in the chick embryo (Bridglal et al., 2021), but that the effects of immobilisation do not become pronounced until day eight and later (Nowlan et al., 2014). Also in the chick embryo, the authors showed that a single day of immobilisation at embryonic days three or four had pronounced, lasting effects on spinal curvature, vertebral segmentation and vertebral shape, while single-day immobilisation at embryonic day five led to the most severe rib abnormalities (Levillain et al., 2019). Drachman and Coulombre (1962) reported relatively severe effects at hatching on the skeleton after one or two days of paralysis between seven and nine days of incubation, indicating that the effects of temporary paralysis (at least in the chick) may become more severe as development progresses. Most published works with a temporal component use the chick or fish model system, and only a small number of studies using mammalian model systems have assessed the effects of absent or abnormal muscle on skeletal development over time (Kahn et al., 2009; Pierantoni et al., 2021). Kahn et al. (2009) assessed joint development in mice with absent or non-contractile skeletal muscle at multiple developmental stages (embryonic day (e)12.5, 13.5, 14.5, 16.5 and 18.5), and reported that elbow joint fusion, first detectable at e16.5 (ca. TS25) is maintained until e18.5 (equivalent to TS27). Joint shape was not quantitatively assessed. Pierantoni et al (2021) quantified humeral bone properties including bone volume and extent at TS23, TS24 and TS27 in muscleless limb and control embryos, and report that while many mineralisation parameters were significantly different at TS24, mineralisation had caught up in the muscleless limb humeri by TS27. Temporal characterisation on the effects of absent muscle on rudiments other than the humerus are lacking, and the progression of joint shape in the absence of skeletal muscle has not been quantified. Given the known differential effects of absent or abnormal muscles on different skeletal rudiments (Kahn et al., 2009; Nowlan et al., 2010), a holistic study of the effects of absent muscle on all key aspects of the developing mammalian skeleton is warranted.

In this work, the progression of prenatal skeletal development when skeletal muscles are absent was investigated, to determine if the effects on the bones and joints remained constant, worsen, or improve over developmental time. Understanding the temporal effects of inter-uterine immobility has impact for conditions such as amyoplasia and developmental dysplasia of the hip, in which an early, often temporary, period of restricted or reduced movement can have long-lasting consequences on joint shape (Nowlan, 2015). In this research, key parameters of skeletal development were assessed, namely joint size, shape and cavitation, and rudiment length and mineralisation, for multiple rudiments and over two developmental stages; TS24 and TS27, in the splotch-delayed “muscleless limb” mouse model and littermate controls. TS24 (e15.5) supplemented our prior studies on TS23 (Nowlan et al., 2010; Sotiriou et al., 2019), while TS27 was the latest prenatal stage which can reliably be obtained in our mouse model. 3D data was obtained using optical projection tomography (OPT), followed by rigid image registration, to characterise morphology, and histology was used for assessment of cavitation.

## Methods

### Animal model

All animal experiments were performed in accordance with European legislation (Directive 2010/63/ EU). Embryos from the Pax3^Sp/Sp^ Splotch delayed (Sp^d^) mice were studied. In Pax3 mutants, muscle progenitor cells do not migrate to the limb buds and thus, the limbs are devoid of skeletal muscle (Franz et al., 1993). Homozygous mutations in Pax3 are neonatal lethal, while heterozygous embryos have no limb muscle abnormalities (Franz et al., 1993). Heterozygous adult animals were imported from The Jackson Laboratory (Maine, USA; JAX stock #000565) and interbred. Pregnant mice were humanely sacrificed using cervical dislocation and embryos were euthanised and staged according to Theiler’s Staging criteria (Theiler, 1989). Genotyping was done by PCR on DNA derived from head tissue. The PCR reaction was carried out for 30 cycles, each with a duration of 30 secconds at 94 °C, 60 °C and 74°C, using three primers. Primer sequences used were AGGGCCGAGTCAACCAGCACG & CACGCGAAGCTGGCGAGAAATG for controls and AGTGTCCACCCCTCTTGGCCTCGGCCGAGTCAACCAGGTCC & CACGCGAAGCTGGCGAGAAATG for mutants. Five muscleless and littermate control embryos at TS24 and at TS27 were analysed in detail in this study. TS24 (around e15.5 in our hands) was chosen as a stage distinct from TS23 which has already been characterised in detail (Nowlan et al., 2010; Sotiriou et al., 2019), and TS27 (around e18.5) was chosen as it is the latest reliably obtainable stage the embryos reach just before birth.

### 3D imaging and image registration

Left fore- and hind-limbs were dehydrated and stained for cartilage and mineralised tissue using alcian blue and alizarin red as previously described (Quintana and Sharpe, 2011a). Stained and fixed limbs were embedded in agarose, dehydrated and cleared in a solution of benzyl alcohol and benzyl benzoate (BABB) in preparation for 3D imaging with optical projection tomography (OPT) following prescribed protocols (Quintana and Sharpe, 2011b). Limbs were scanned under visible light to obtain 3D images of the alcian blue staining (cartilage) and under the Texas-red filter to obtain auto-fluorescent 3D images of the alizarin red stained region (the mineralised region). Scans were reconstructed using NRecon (SkyScan, Brucker microCT, 2011). Segmentation of the cartilaginous scapula, humerus, ulna, radius, pelvis, femur and tibia was performed with Mimics (v17.0, Materialise, Leuven, Belgium). The fibula was prone to deformation and twisting, making it difficult to reliably segment and was therefore not included in any analyses. As previously described (Sotiriou et al., 2019), segmentation was still possible when rudiments were fused, due to the joint line being still identifiable based on reduced intensity of staining. Segmented rudiments were prepared for image registration using TransformJ (Meijering et al., 2001) in ImageJ (Schneider et al., 2012), being first roughly aligned with other rudiments of the same type, scaled to a consistent magnification, and allocated the same canvas area. Image brightness was normalised using Matlab (version R2015a, the MathWorks, Inc., Massachusetts, USA). The Image Registration Toolkit software (Schnabel et al., 2001) (IRTK, BioMedia, Imperial College London) was used to align each rudiment to each other, with rigid registration as previously described (Sotiriou et al., 2019). Segmentation or image registration were not necessary for the alizarin red scans, due to the simple nature of the bone collar measurements taken.

### Measurements and statistics

For joint shape characterisation, the same set of 25 forelimb measurements and 27 hindlimb measurements described in our previous study of muscleless limb joint shapes at TS23 were used, reproduced in Figures 1 and 2 (with permission). Consistent measurements were made from equivalent sections and planes of each individual rudiment by applying the same rotations to each registered rudiment dataset. Measurements were performed in Gwyddion image editing software (Gwyddion 2.44, http://gwyddion.net/, last accessed September 2021). As differences in rudiment length were previously reported for some muscleless rudiments (Nowlan et al., 2010) measurements were normalized by the length of the rudiment under investigation, in order to focus the outcomes on shape-specific (rather than overall size dependent) changes.

**Figure 1.**
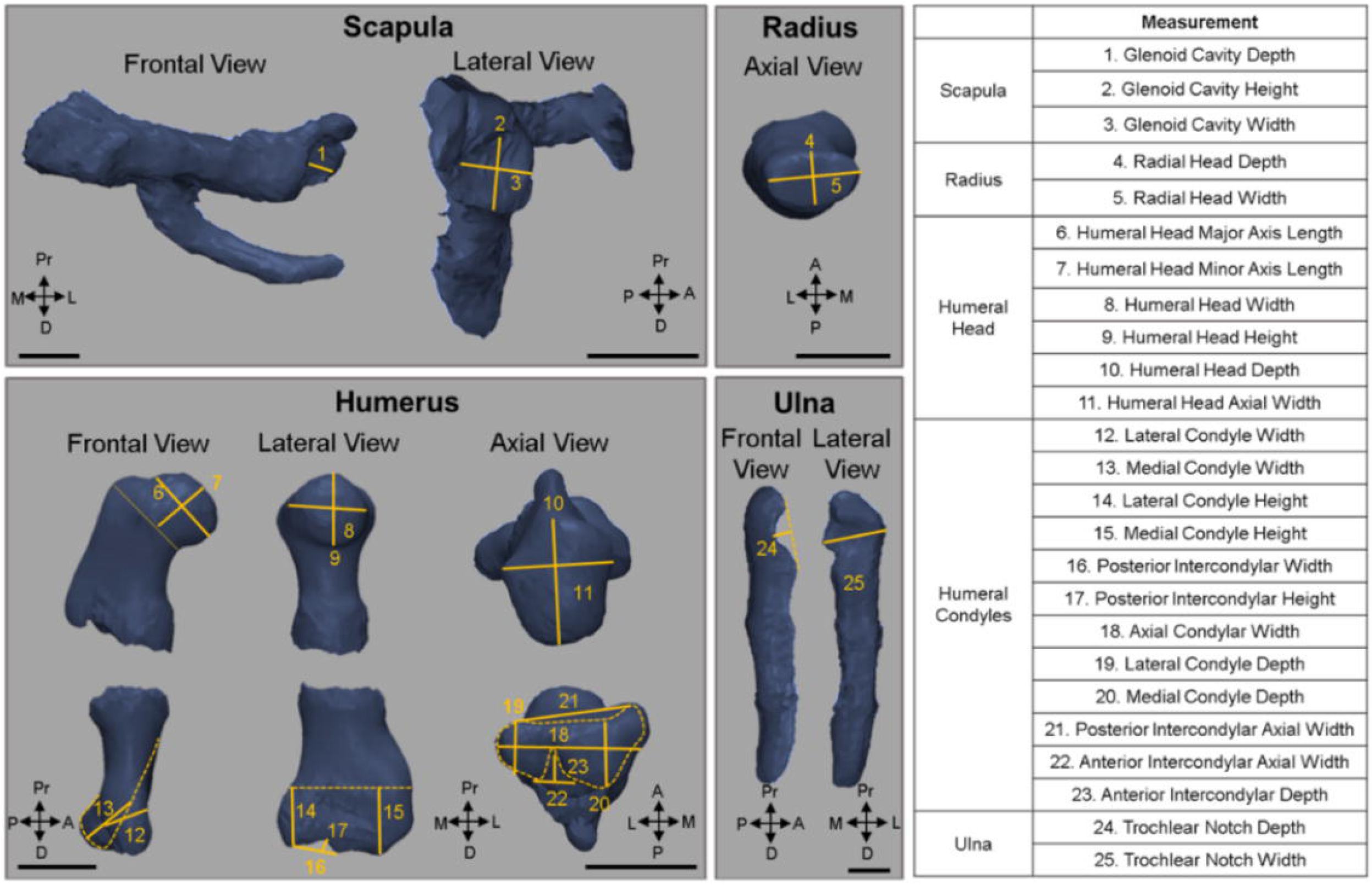
Measurements made on forelimb joints. Image reproduced from [10] © 2019 Orthopaedic Research Society. Published by Wiley Periodicals, Inc.

**Figure 2.**
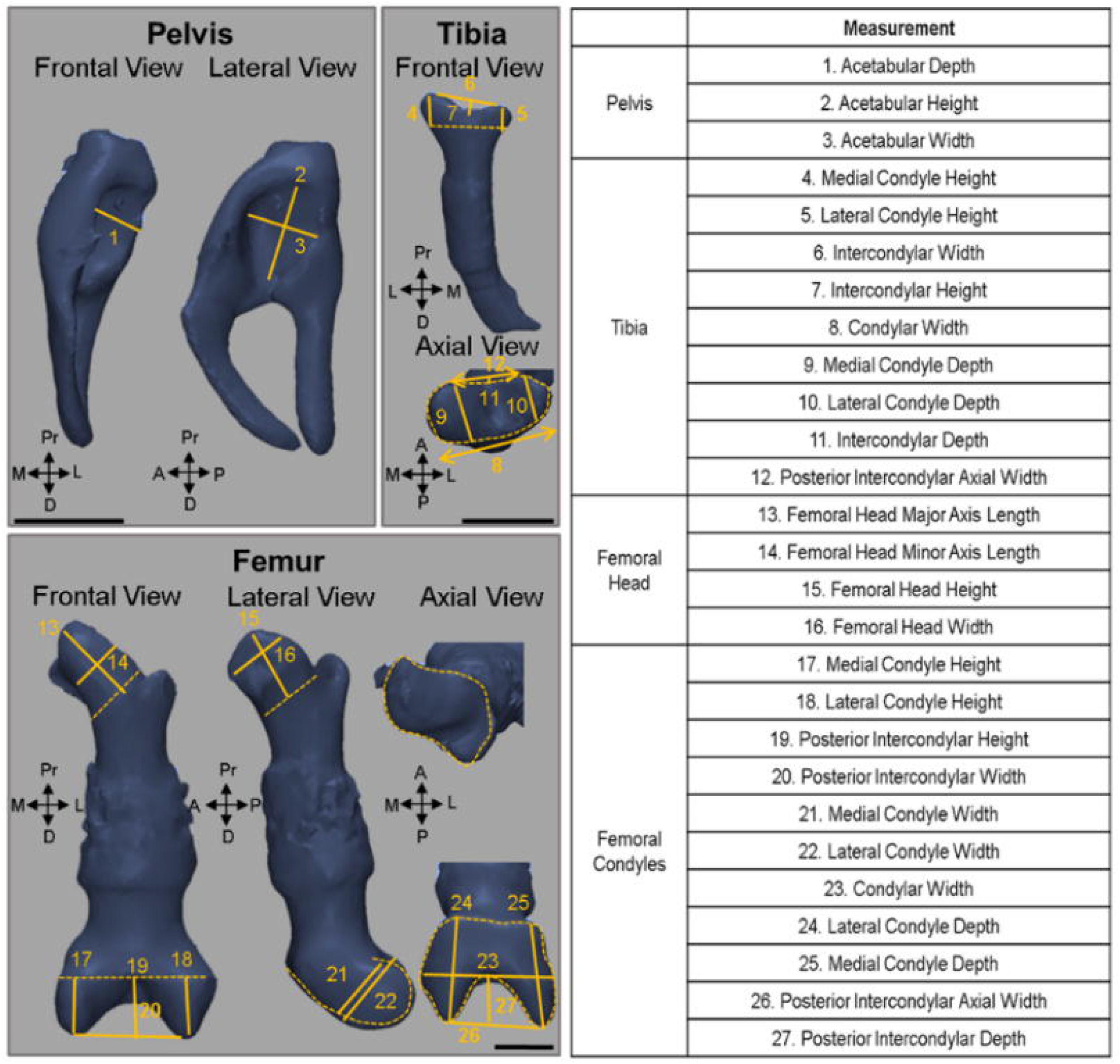
Measurements made on hindlimb joints. Image reproduced from [10] © 2019 Orthopaedic Research Society. Published by Wiley Periodicals, Inc.

Rudiment lengths were measured on a lateral view. Mineralisation of the long bones was measured on a frontal cross section of the alizarin red scans. In cases where the mineralisation extent was uneven between the medial and lateral aspects, the average of the two extents was taken. Mineralisation of the pelvis and scapula was not quantified, due to difficulties in consistent measurement of the mineralisation extent in flat bones. Long bone rudiment lengths, absolute mineralisation extent, and mineralisation extent normalised to rudiment length were presented graphically.

All datasets were tested for normality with the Shapiro-Wilk test. For datasets with a normal distribution, two tailed Student’s t tests for independent samples (SPSS Statistics 24, IBM corp., Armonk, NY) were performed to determine which measurements were statistically significant between control and muscleless groups. For non-normal datasets, Mann-Whitney tests were performed. Joint shape measurements for which a significant difference (p < 0.05) between controls and muscleless limb mutants were found were displayed graphically, while the full table of results is available on Zenodo (https://doi.org/10.5281/zenodo.5566902). To allow for visual assessment of changes in shape, rudiment shape outlines were traced on frontal, lateral, and axial sections through the prime regions of interest for each rudiment.

### Histology

Cavitation of the glenohumeral (shoulder), elbow, hip and knee joints was assessed in the control and the muscleless limb embryos using standard histology. Limbs were dissected and processed for cryo-sectioning in an increased sucrose gradient (15% and 30% sucrose) as described previously (Ahmed and Nowlan, 2020). Processed limbs were embedded in OCT (optimal cutting temperature) (Agar Scientific, Stansted, UK) diluted with 50% sucrose and cut (12 μm thickness) using a cryostat (NX70, Leica Biosystems, UK). Then, frozen sections were fixed with 4% (w/v) paraformaldehyde, stained with 0.1% toluidine blue (Sigma-Aldrich) for 3 s and washed with tap water. Following air-drying, sections were imaged by transmitted illumination using a light microscope (Yenway EX30; Life Sciences Microscope, Glasgow, UK).

## Results

### TS24 Forelimb

#### Glenohumeral Joint

The muscleless limb glenoid cavity at TS24 was abnormally shaped in the lateral plane when compared to the glenoid cavity of the controls. The outlines of the glenoid cavity of the muscleless limb scapulae were more elliptical than the controls and exhibited an ectopic protrusion distally (Figure 3B, black arrow). The height and width of the glenoid cavity in the muscleless limb embryos were significantly larger than controls (Figure 3, measurements 2 and 3 respectively). Further shape abnormalities were present in the proximal humerus. The muscleless limbs had a visibly elongated humeral head in the frontal plane compared to controls (Figure 3C, purple outlines) which was reflected in a significant decrease in the major axis length of the humeral head (Figure 3, measurement 6). On the lateral plane, the shape of the muscleless limb humeri was abnormal resembling an ellipse with a tilted major axis (Figure 3D, dotted line). The glenohumeral joints of the TS24 muscleless limb embryos were fused with no clear separation between the glenoid cavity of the scapula and the humeral head (Figure 4, filled arrowhead).

**Figure 3.**
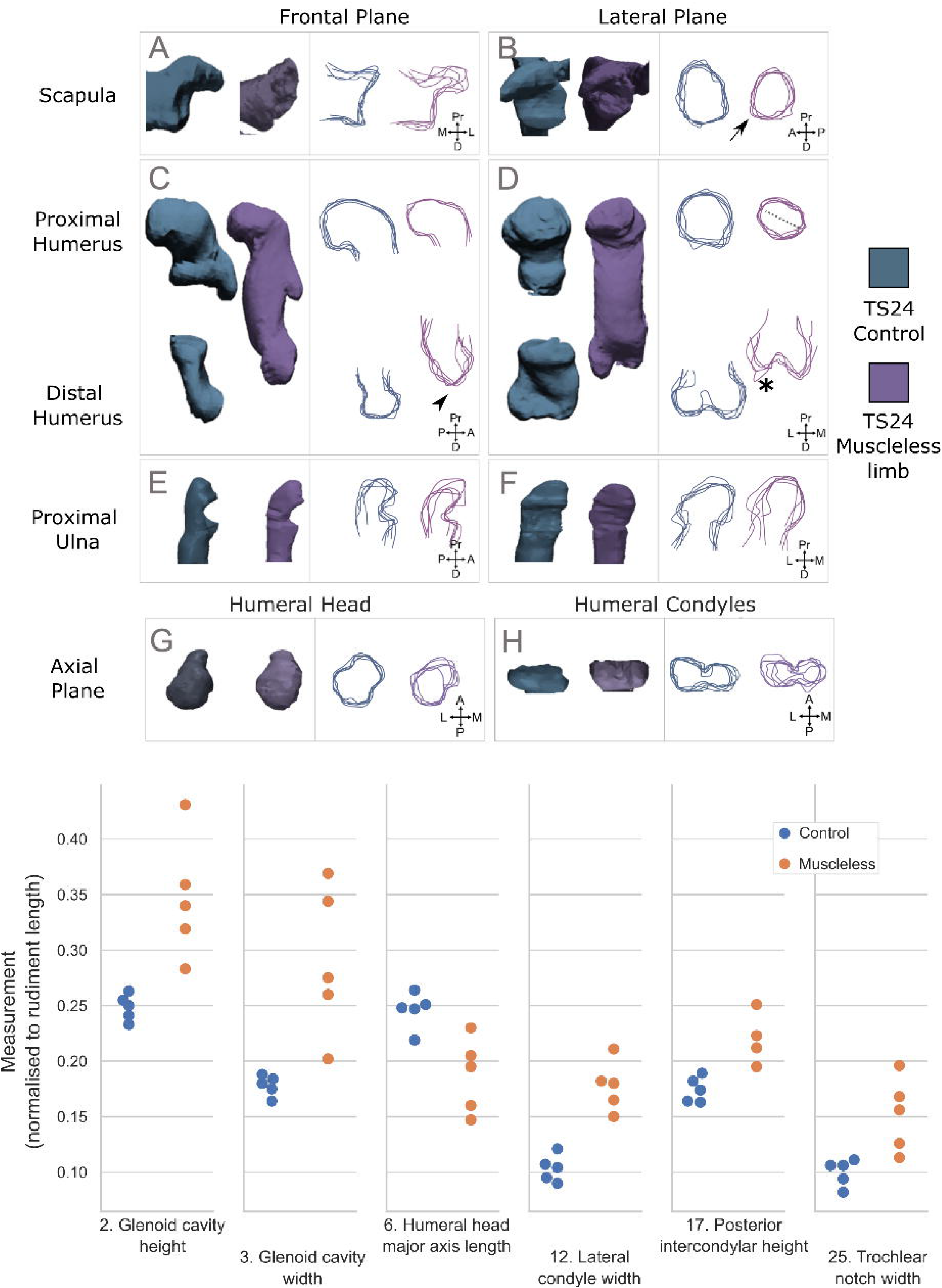
Qualitative and quantitative joint shape characterisation of the forelimb joints at TS24. Above: Representative 3D shapes and outlines of the scapula, humerus, radius and ulna of control (blue) and muscleless limb (purple) embryos at developmental stage TS24. Arrow in B indicates abnormal protrusion of muscleless glenoid cavity, arrowhead in C represents abnormal shape of distal humerus and asterisk in D indicates abnormal shape of lateral condyle in distal humerus of muscleless limbs. Below: Dot plots illustrating all hindlimb measurements with significant differences (p-value<0.05) between muscleless limb (orange) and control (blue) groups.

**Figure 4.**
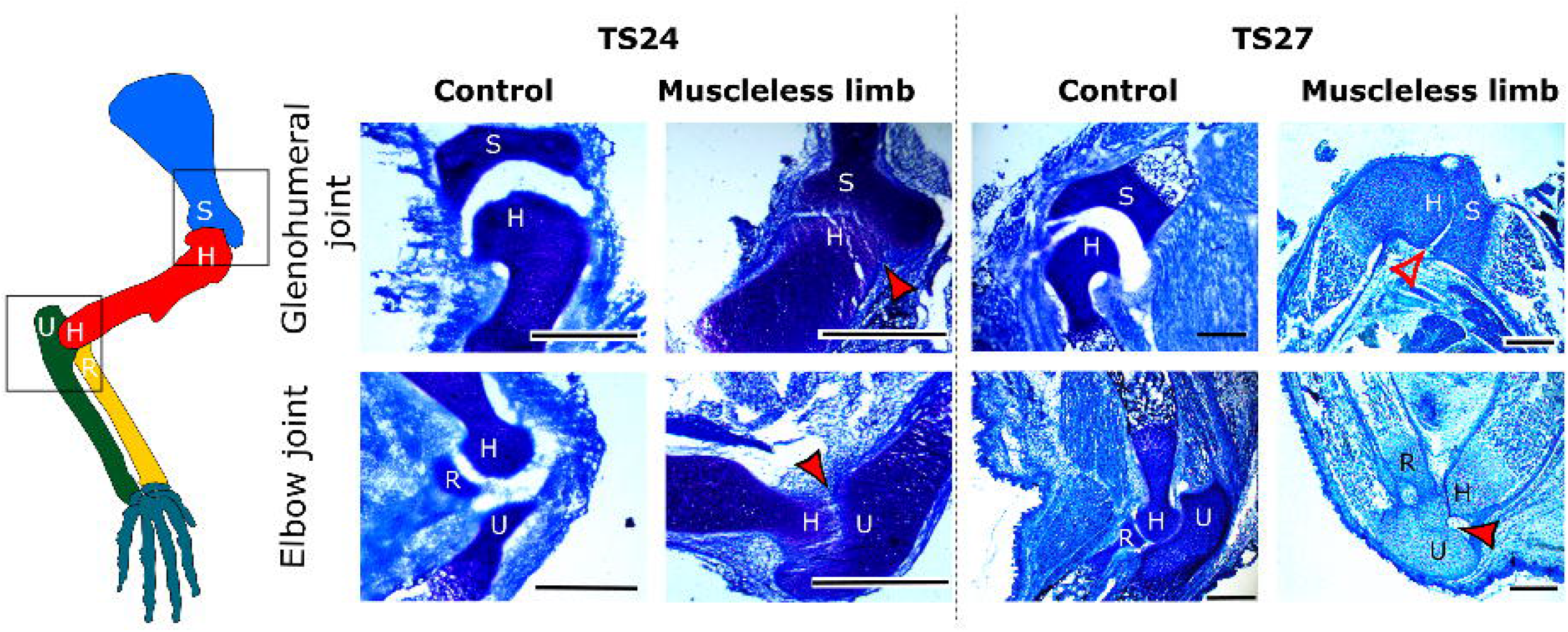
Cavitation in the forelimb joints at TS24 and TS27. S: scapula, H: humerus, U: ulna, R: radius. The glenohumeral joints at TS24 were not cavitated (filled arrowheads), with partial cavitation at TS27 (hollow arrowhead). The elbow joints were fused at both TS24 and TS27 (filled arrowheads). Representative images shown-results were consistent for all three samples per stage. Scale bars 500 μm.

#### Elbow Joint

The lateral condyle of the TS24 muscleless limb humerus had an abnormal, angular protrusion on the lateral plane (Figure 3D, asterisk) a result corroborated by the significantly greater posterior intercondylar height (Figure 3, measurement 17). The lateral condyle of muscleless limbs was also significantly wider than controls (Figure 3, measurement 12). The medial condyle of the muscleless limb humerus was irregular in the frontal plane with its shape not having the bulbous shape of the controls (Figure 3C, arrowhead). The shape of the muscleless limb distal humerus in the axial plane varied a lot from sample to sample, whereas that of the controls was more consistent between samples (Figure 3H, outlines). In the rudiments opposing the distal humerus, the muscleless limb ulnae were less intricately shaped in the lateral plane than the ulnae of the control embryos (Figure 3F, outlines) and the trochlear notch was significantly wider (Figure 3, measurement 25). The radii of the muscleless limb embryos were not different in shape to the controls and no significant differences in the shape of the proximal radii were found. Histologically, the elbow joints were fused at TS24 with no separation between the humeral condyles and the radius and ulna (Figure 4, filled arrowhead).

### TS27 Forelimb

#### Glenohumeral Joint

At TS27, the muscleless glenohumeral joints were not as pronouncedly different in qualitative shape to the controls as at TS24. No significant differences in the TS27 glenoid cavity were detected. The glenoid cavity was elliptically shaped in both control and muscleless limb groups and only one out of five muscleless limb scapulae still exhibited an abnormal protrusion at the lower end of the long axis (Figure 5B, arrow). The humeral head of the muscleless limb embryos resembled that of the controls in the frontal plane (Figure 5C), but in the lateral plane, the muscleless humeral heads were visibly and quantitatively wider compared to the controls (Figure 5D; dotted line & Figure 5; measurement 8). In the axial plane (Figure 5G), the muscleless limb humeri had an indentation on the antero-lateral aspect (arrowhead) which was not present in the controls, and malformation of the humeral head was also detected though a significant difference in the humeral head axial width (Figure 5, measurement 10). Therefore, at TS27, the proximal humerus was more severely affected by the lack of muscle than the glenoid cavity. At TS27, cavitation at the glenohumeral joint had consistently occurred in the muscleless limb embryos (Figure 4B), but with a less prominent separation between the scapula and humeral head as seen in the controls (Figure 4, hollow arrowhead).

**Figure 5.**
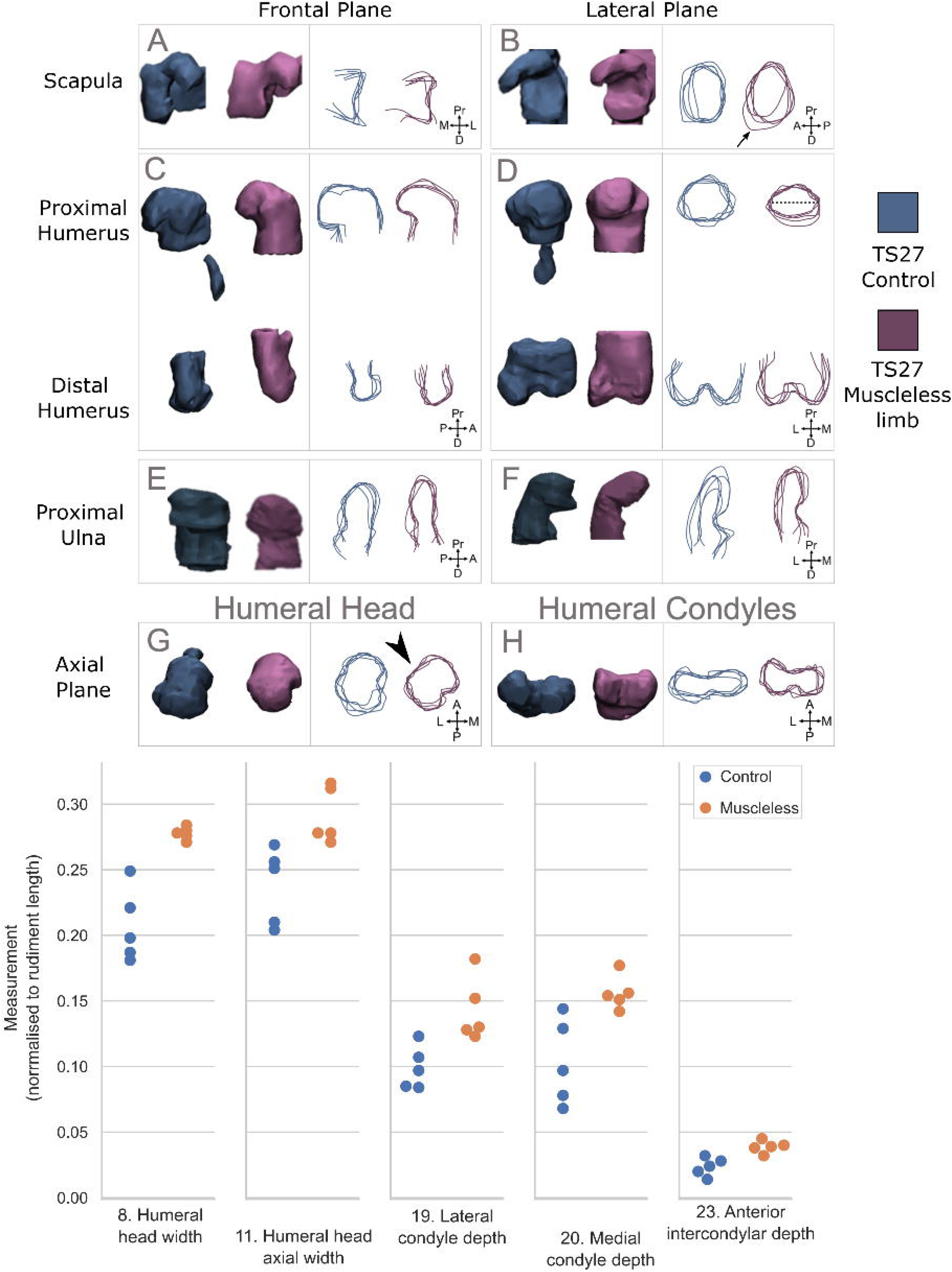
Qualitative and quantitative joint shape characterisation of the forelimb joints at TS27. Above: Representative 3D shapes and outlines of the scapula, humerus, radius and ulna of control (blue) and muscleless limb (purple) embryos at developmental stage TS27. Dashed line in D indicates wider appearance of humeral head in the lateral plane. Arrowhead in G indicates the abnormal shape of the muscleless limb humeral head in the axial plane. Below: Dot plots showcasing the significant differences (p-value<0.05) in measurements between the muscleless limb (orange) and control (blue) groups for the rudiments comprising i) the glenohumeral joint, and ii) the elbow joint.

#### Elbow Joint

The shapes of the distal humerus, and proximal radius and ulna at TS27 were qualitatively similar in both control and muscleless limb embryos in the frontal and lateral planes (Figure 5). However, in the axial plane, the muscleless humeral condyles appeared different to those of controls (Figure 5H), which was reflected in three significantly different measurements all made on the axial plane, namely significantly increased lateral, medial condyle and anterior intercondylar depths in the muscleless limbs compared to the controls (Figure 5, measurements 19, 20 and 23 respectively). There were no significant differences in the proximal ulna between the muscleless limb and control groups at TS27. As at TS24, no shape or size differences were observed between the radii of the two groups at TS27. The elbow joint continued to be fused at TS27 (Figure 4, filled arrowhead). Therefore, a partial recovery of elbow joint shape occurred in the muscleless limb forelimb joints by TS27, despite the lack of cavitation.

### TS24 Hindlimb

#### Hip Joint

At TS24, the acetabulum was qualitatively and quantitatively deeper and wider in the muscleless limb group than the controls (Figure 6 A, measurements 1 & 3). The shape of the muscleless acetabulum was also more variable in the lateral plane than in controls (Figure 6B). The muscleless limb femoral head appeared longer and more slender than in controls (Figure 6C, D) and the femoral head height was significantly greater in the muscleless limbs than in controls (Figure 6, measurement 15). These features in the muscleless proximal femur may indicate an adjustment of femoral head shape to the deeper concavity of the acetabulum in this group. At TS24, the hip joint of control limbs was clearly separated (Figure 7), while the muscleless limb hip was fused with no separation between the femoral head and the acetabulum (Figure 7, arrowhead).

**Figure 6.**
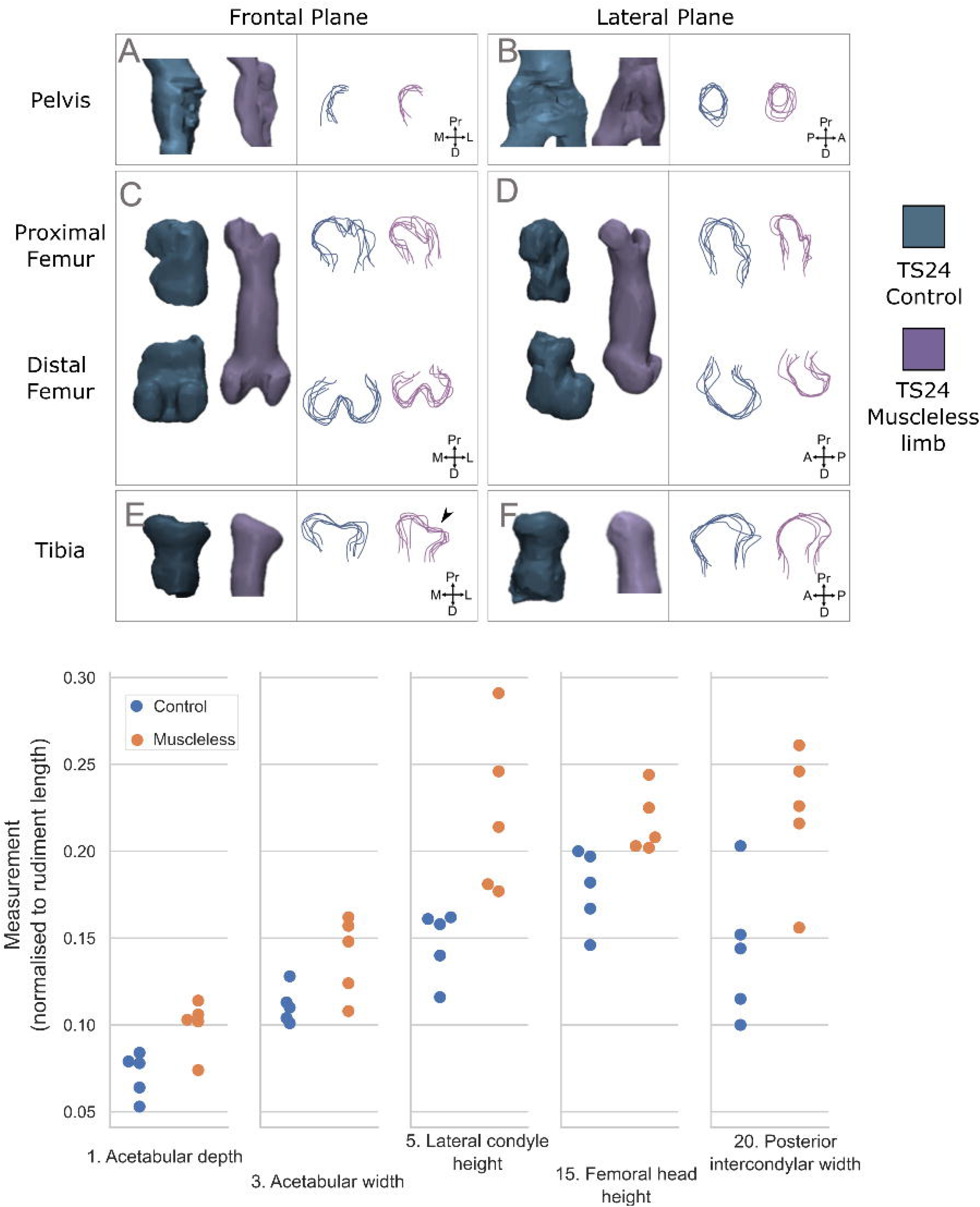
Qualitative and quantitative joint shape characterisation of the hindlimb joints at TS24. Above: Representative 3D shapes and outlines of the pelvis, femur and tibia of control (dark blue) and muscleless limb (dark purple) embryos at developmental stage TS24. Arrowhead in E indicates abnormal shape of proximal tibia. Below: Dot plots illustrating all hindlimb measurements with significant differences (p-value<0.05) between muscleless limb (orange) and control (blue) groups.

**Figure 7.**
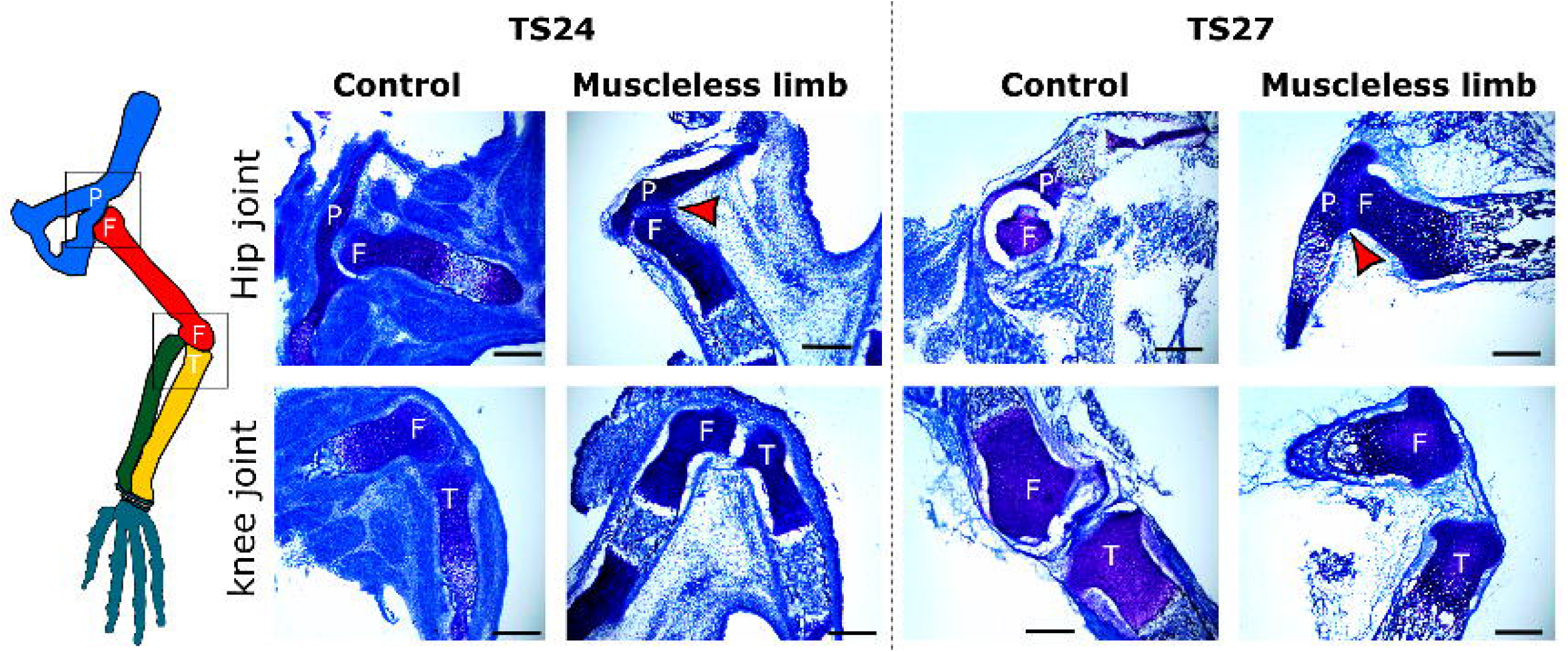
Cavitation in the major hindlimb joints at TS24 and TS27. P: pelvis, F: femur, T: tibia. Cavitation was absent in the hip joint at TS24 and TS27, and present in the knee joint at both stages. Representative images shown-results were consistent for all three samples per group/stage. Scale bars 500 μm.

#### Knee joint

On the frontal plane, both muscleless limb femoral condyles at TS24 had a more angular shape when compared to the condyles of the controls with the intercondylar region resembling a triangle rather than a curve (Figure 6C), reflected in a significant difference in the posterior intercondylar width (Figure 6, measurement 20). The lateral condyle of the muscleless limb tibiae protruded more prominently out from the diaphysis than controls (Figure 6E, arrowhead), reflected in a significant increase in the tibial lateral condyle height in the muscleless limb group compared to controls (Figure 6, measurement 5). Knee joints of the muscleless limb embryos was fully cavitated at TS24, as was the knee joint of the control embryos at the same stage (Figure 7).

### TS27 Hindlimb

Remarkably, both the hip and the knee joints of the muscleless limb embryos at TS27 resembled in shape those of the controls (Figure 8). None of the obvious shape differences seen at TS24 remained. This was corroborated by the lack of significant differences in any of the measurements performed on the articular surfaces comprising the two joints at TS27. The hip joint of the TS27 muscleless limb embryos remained fused as no separation was seen between the femur and the acetabulum (Figure 7, arrowhead), while the muscleless limb knee joint remained fully cavitated (Figure 7). Therefore, in muscleless limb embryos at TS27 both hip and knee joint shape recovered so as to be equivalent to control shapes, despite the continued lack of cavitation in the hip joint.

**Figure 8.**
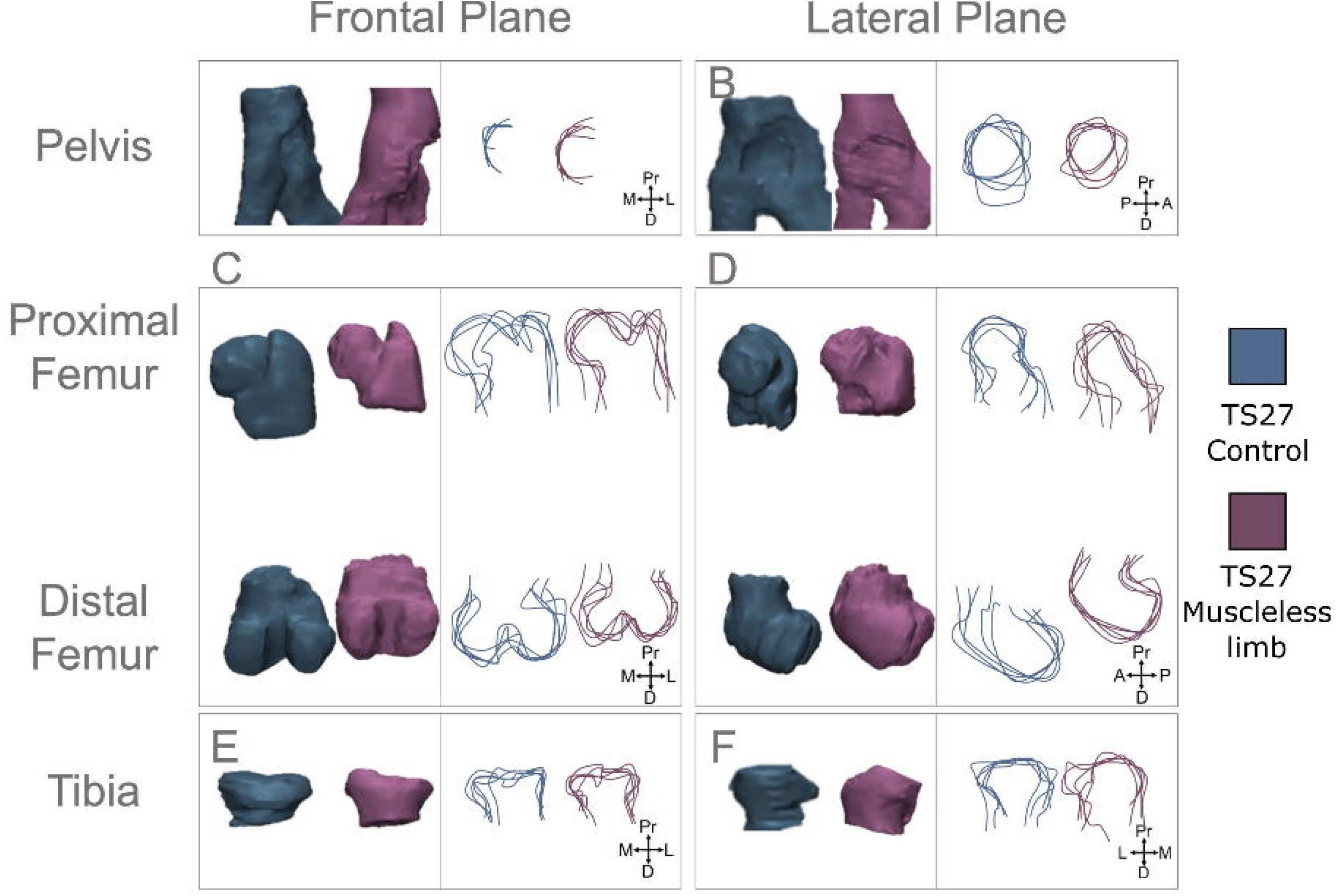
Qualitative joint shape characterisation of control (blue) and muscleless limb (purple) hindlimb joints at TS27. No measurements of the hindlimb joint shapes showed any significant differences at TS27.

### Rudiment Length and Mineralisation

#### Forelimb

At TS24, the humerus, ulna and radius were all significantly shorter than controls of the same stage (Figure 9). At TS24, these three rudiments also had significantly reduced mineralisation in muscleless limbs compared to controls, for both absolute mineralisation extent, and adjusted for rudiment length (Figure 9). At TS27, the muscleless humerus and ulna were still significantly shorter than controls, while there was no significant difference between the muscleless and control radii (Figure 9). A dramatic change in mineralisation progress had occurred by TS27. Mineralisation of the three forelimb muscleless rudiments “caught up” with controls of the same age, with no significant reductions in absolute or length-proportionate mineralisation in any of the three rudiments. In fact, the proportion of mineralisation in the muscleless ulna at TS27 actually significantly exceeded that of the control ulna (Figure 9).

**Figure 9.**
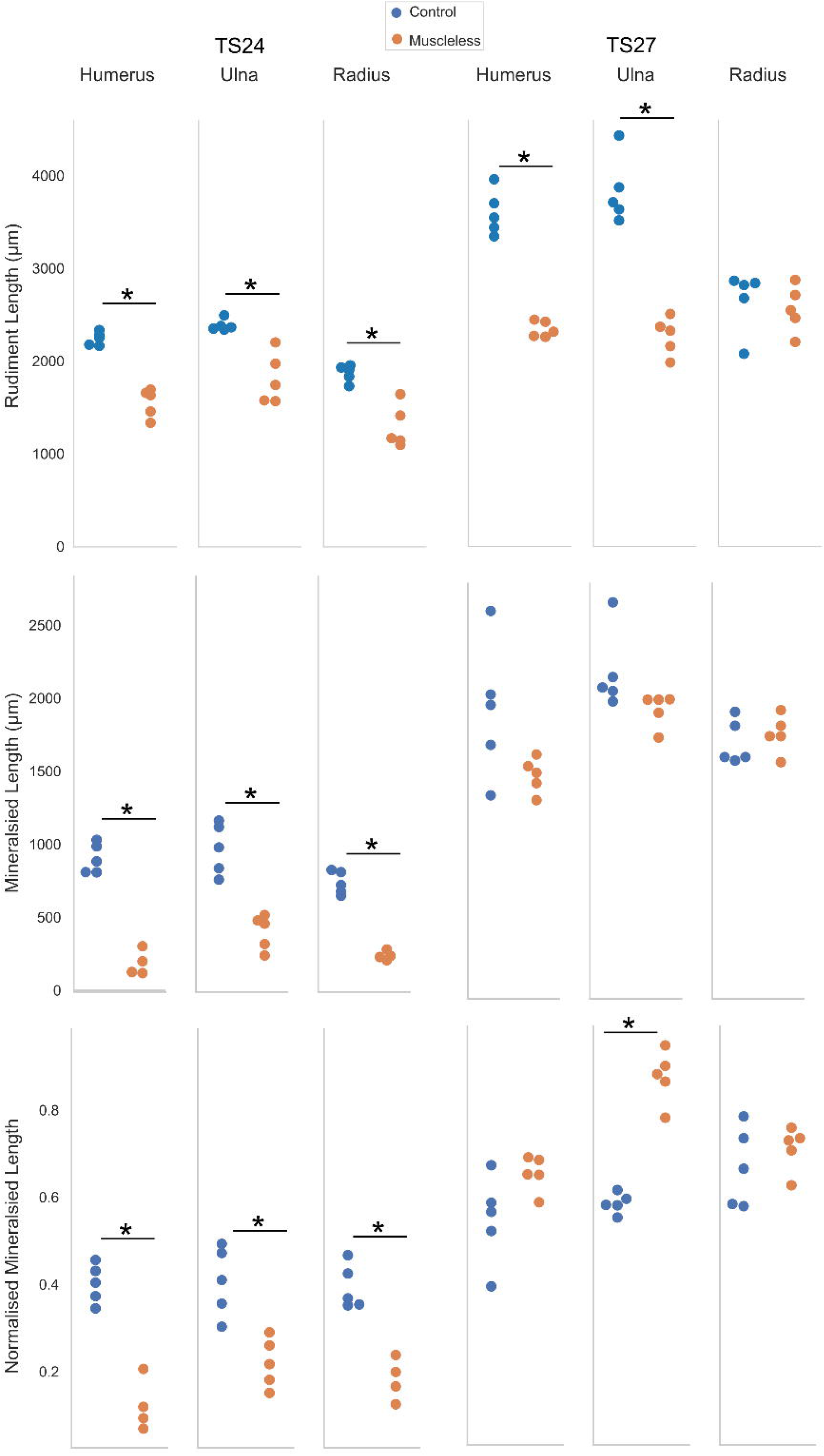
Forelimb rudiment length and mineralisation. At TS24, all length and mineralisation measurements were significantly reduced in muscleless forelimbs (orange) compared to controls (blue). At TS27, humeri and ulnae were still significantly smaller in the muscleless limbs, while all mineralisation extent measures had recovered, or even exceeded, those of littermate controls. Normalised mineralisation lengths calculated by mineralised length divided by rudiment length (equivalent to mineralised proportion). *p<0.05

#### Hindlimb

At TS24, both the femur and tibia were significantly shorter in the muscleless limbs than in the controls, and the mineralisation extent of these rudiments (absolute and length-normalised) were also significantly reduced in the muscleless limbs compared to the controls (Figure 10). By TS27, there was no longer a significant difference in femoral length between the groups, while the tibia was still significantly shorter in the muscleless limbs than in the controls (Figure 10). As in the TS27 forelimb, mineralisation of the muscleless hindlimb rudiments had recovered with respect to controls of the same age, with no significant reductions in absolute or length-proportionate mineralisation relative to controls (Figure 10). As with the ulna, mineralisation of the tibia (adjusted for rudiment length) actually exceeded that of the control tibiae (Figure 10).

**Figure 10.**
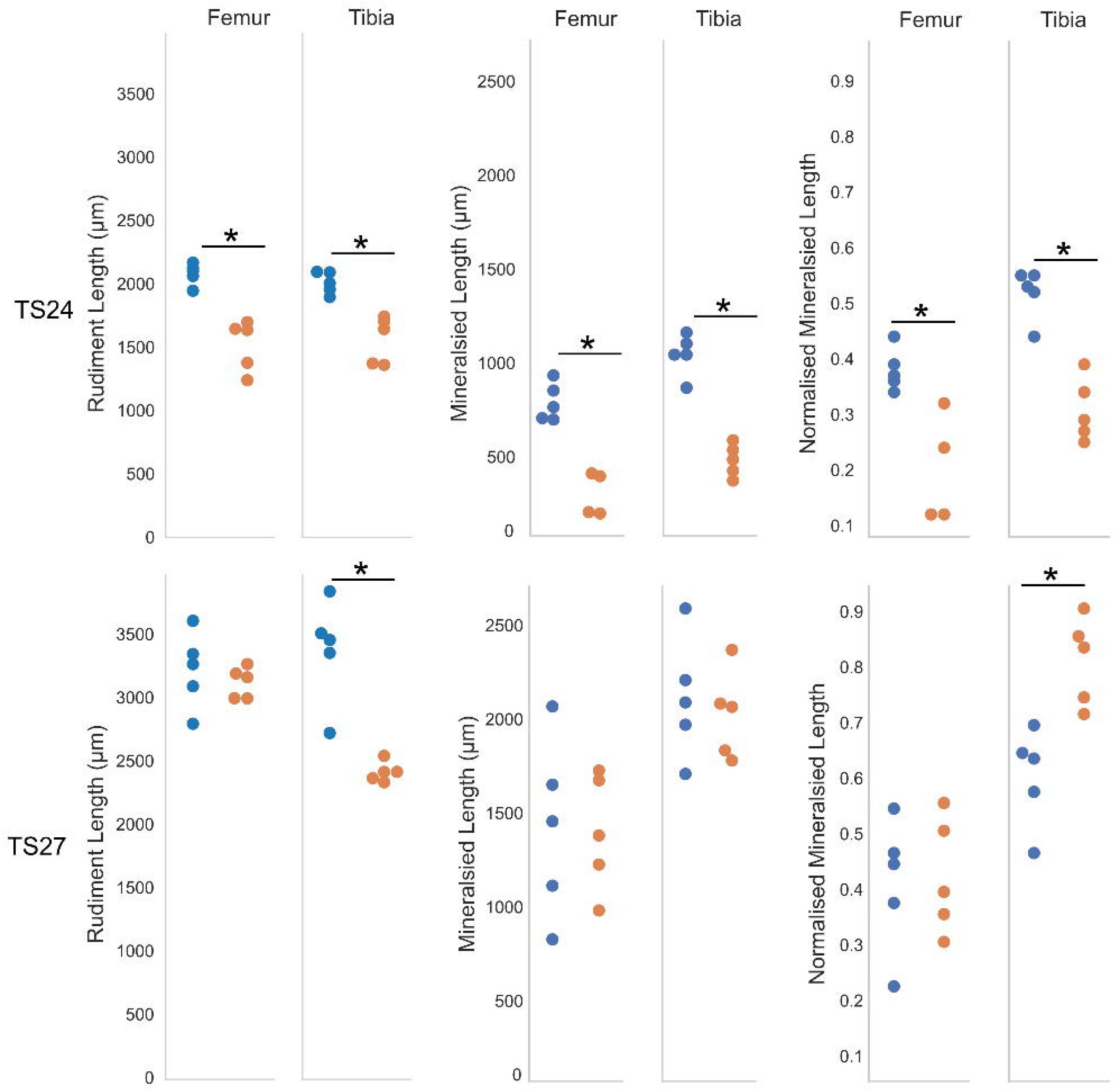
Hindlimb rudiment length and mineralisation. At TS24, femoral and tibial length and mineralisation measures were significantly reduced in muscleless limbs (orange) compared to controls (blue). At TS27, tibial lengths were still significantly lower in the muscleless limbs, while mineralisation extent measures had recovered, or even exceeded, those of littermate controls. Normalised mineralisation lengths calculated by mineralised length divided by rudiment length (equivalent to mineralised proportion). *p<0.05

## Discussion

The effects of absent skeletal musculature on the progression of joint shape and cavitation, rudiment length and rudiment mineralisation in the mouse embryo were assessed. By TS27 (prenatal stage), hip and knee joint shape had recovered. Partial recovery of glenohumeral and elbow shape in muscleless limbs at TS27 was observed, with some rudiments having recovered more than others. Cavitation did not follow the same pattern as joint shape in the muscleless limbs as development progressed. Joints which had not cavitated at TS24 remained completely or almost completely uncavitated at TS27. Only the glenohumeral joint showed an improvement in cavitation status with advancing developmental age. In contrast, mineralisation of the TS27 muscleless limb long bone rudiments caught up with- and in some rudiments even exceeded-mineralisation in controls.

Although active muscle contractions are completely absent in the limbs of the Splotch-delayed mouse embryos, there are still two sources of mechanical stimulation acting on or in the rudiments. The first is mechanical stimulation from passive movements due to the activity of the mother and the normal littermates. The authors previously demonstrated that 10 μm passive movements induce differential levels of biophysical stimuli between the forelimb and hindlimb at TS23 and TS24, and proposed that such movements contribute to the more pronounced effects of absent muscle on the forelimb than on the hindlimb (Nowlan et al., 2012). While passive movements would have a negligible effect in normal embryos (due to the far larger active movements induced by muscle contractions (Nowlan et al., 2012)), it is reasonable to believe that they are playing a role in the recovery of aspects of skeletal development in the muscleless limb mice. As pregnancy progresses, the space each embryo has decreases, which could increase the intensity and impact of passive stimuli in the muscleless limbs, potentially contributing to the recovery of joint shape, rudiment growth and rudiment mineralisation. The other source of mechanical stimulation acting during development, including in the muscleless limbs, is growth-generated strains and pressures. First proposed as acting like a complex field of morphogens influencing early limb morphogenesis in 2002 by Henderson and Carter, growth-generated strains and pressures arise during development when *“complex configurations of connected tissues grow at different rates”,* leading to *“the generation of time-varying, quasi-static stresses and strains throughout the developing cells and tissues”* (Henderson and Carter, 2002). During prenatal development with normal skeletal muscle and movements, growth-related strains and pressures likely play a very minor role, once movements have commenced. However, when skeletal muscle is absent, the biophysical stimuli arising from growth-related strains and pressures could contribute to growth, morphogenesis and differentiation in the muscleless limb skeleton.

Our findings that forelimb joint shape and cavitation, and rudiment length and mineralisation were more severely affected by a lack of skeletal muscle than the hindlimb joints and rudiments confirmed prior reports (Kahn et al., 2009; Nowlan et al., 2010). Our findings that the effects of absent muscle on mineralisation reduce as development progresses also confirms prior work from the authors (Pierantoni et al., 2021). The advance provided by the current work is it is the first demonstration or partial or full recovery of joint shape, rudiment length and rudiment mineralisation from the effects of absent skeletal muscle by the latest prenatal stage. Intriguingly, cavitation was the only aspect not to recover to some extent over development. This may indicate that cavitation is a single “rupture” type event, as previously proposed (Drachman and Sokoloff, 1966), and if movement does not occur at the critical time, the opportunity for normal cavitation is lost. It is feasible that active bending movements, localised to the joint, are needed for cavitation to occur, which would not occur even with increasing growth-generated stresses or strains due to passive movements. This concurs with our recent study, in which a cavity was artificially induced in the developing chick using an applied bending movement (Bridglal et al., 2021). What was surprising was the partial or complete recovery in joint shape despite the lack of, or incomplete, cavitation in the glenohumeral, elbow and hip joints, which contradicts our prior understanding of joint shape being heavily dependent on successful cavitation (Bridglal et al., 2021). Another interesting feature of the joint shape results was that, in the glenohumeral and elbow joints, different features were abnormal at TS27 than at TS24. This could imply that, as the joints continue to grow, the opposing surfaces are continually shaping and moulding each other, providing the opportunity for some internal correction of early abnormalities. Shape abnormalities newly arising at TS27 could be due to some form of compensation for early shape abnormalities in the opposing surface, or alternatively, particularly in the elbow joint, could be due to the continued joint fusion. However, as the hip joint did not have any quantitative shape differences at TS27, despite the ongoing lack of cavitation, this would indicate that perhaps local interaction between opposing joint surfaces is the primary driver of shape recovery, rather than the capacity for movement at the joint.

One of the most fascinating findings was the remarkable recovery of mineralisation extent in the muscleless limbs between TS24 and TS27, with two rudiments (the ulna and tibia) at TS27 actually having significantly more mineralisation (proportionate to rudiment length) compared to controls at the same age. What prompted the dramatic acceleration in mineralisation over the three day period between TS24 and TS27? The initiation and early progression of ossification is evidently delayed in the muscleless limb mice, as previously described (Nowlan et al., 2010). Is this early delay compensated for by accelerated mineralisation after TS24, and if so, what is the mechanism underlying this compensation? Progression of the growth plates and of mineralisation has been shown to be promoted by cyclic hydrostatic pressure (Henstock et al., 2013) or applied compression loading (Khatib et al., 2021) and we theorise that increasing biophysical stimuli due to passive movements (as intrauterine space decreases) and due to growth-generated strains (as the rudiments continue to grow) accelerate the progression of mineralisation in the muscleless limb mice. The question remains as to why the mineralisation proportion in the ulna and tibia was actually higher than in controls at TS27. The muscleless ulnae and tibiae remained significantly smaller in length than controls at TS27, and therefore an accelerated mineralisation rate, operating independently of rudiment length, could have led to the greater mineralisation proportion in the muscleless limbs. The ulna and tibia are both the dominant rudiments in the zeugopod (the middle section of the limb). They would likely be subject to lower biophysical stimuli from passive movements compared to the stylopod (most proximal region of the limb), due to greater bending moments in the stylopod. Another possibility is that the accelerated mineralisation of the zeugopod is due to an increased level of growth-generated strains based on restraint between the strong stylopod rudiment, and the complex, set of joints at the end of the stylopod at the paws, with lateral restraint due to the presence of the radius or fibula. It is worth highlighting that the recovery in mineralisation extent does not necessarily mean a recovery in the ossification process, as the mineral deposited in the muscleless limbs could be abnormal or inferior in quality.

There are some limitations to our research. The first is that the animal numbers are quite small, with a group size of five. However, the results were reasonably consistent, with variation within groups dwarfed by the variation between control and muscleless limb groups, as can be seen from both the raw data (dot plots) and the rudiment outlines. Another limitation is that our method of normalising shape measurements to rudiment length has the potential to disproportionately skew shape features which may not change in proportion to length. However, it was reassuring that significantly different measurements correlated with shape abnormalities visible in the outlines, and that shape recovery at TS27 was evident both in the qualitative and the quantitative analyses. As the Pax3 mutation is neonatal lethal, it is unfortunately not possible to quantify the effects of absent skeletal muscle on a longer timescale than in this research. Despite the dramatic improvement in the various parameters of skeletal development studied by TS27, the skeletons would not be expected to be normally functioning after birth (if, in theory, the mice would survive). The lack of cavitation in particular would inhibit movement and potentially lead to new or worsening joint shape abnormalities. Furthermore, even very slight shape abnormalities not detected by our measurements would likely impact on function and health of the joint. For example, the shape abnormalities involved in hip dysplasia are not always very dramatic, and yet with growth and over time, there are severe consequences for gait and joint health.

The relevance of the current study for human conditions is that multiple aspects of the developing murine skeleton had a remarkable capacity for recovery despite the absence of skeletal muscle and the active movements resulting from muscle contractions. Developmental dysplasia of the hip and amyoplasia result from a period of abnormal or restricted intra-uterine movement (Nowlan, 2015). Hip dysplasia is diagnosed with ultrasound after four postnatal weeks (Tan et al., 2019), and amyoplasia (despite its quite dramatic impact on skeletal morphology) is diagnosed prenatally in only 22.2% of cases (Filges and Hall, 2013). Could it be the case that the incidence of skeletal abnormalities is much greater *in utero*, but that majority of these have self-corrected and resolved prior to birth? Many cases of hip instability in neonates resolve without any intervention (Bialik et al., 1999), and hip dysplasia is likely more a spectrum than a dichotomous diagnosis, where potentially only the most severe cases do not spontaneously recover either pre- or post-natally. The fact that restricted movement postnatally, for example in a cradle board (Dezateux and Rosendahl, 1997), can induce hip dysplasia in formerly healthy hips, further demonstrates the critical role of muscle, movements and mechanics in healthy skeletal development. Given that late or missed diagnoses of hip dysplasia are a significant issue, usually necessitating surgery (Broadhurst et al., 2019), it is also possible that a hip joint which satisfies the screening criteria in early postnatal life could adapt and adjust its shape negatively as the baby grows. A key question which was not answerable in this research is how the ability to self-correct and recover abnormal skeletal development declines over postnatal development. If adaptation potential of the developing skeleton declines over postnatal development, it may make sense to harness the plasticity of the developing skeleton at as young an age as possible, through physical therapy, casting or harnesses. This question will be explored in future work, with an alternative postnatal animal model.

In conclusion, skeletal development was studied at two different prenatal ages in the muscleless limb mouse. With advancing prenatal development, the effects of absent muscle on all parameters, apart from joint cavitation, become less severe. Joint shape, rudiment length, and rudiment mineralisation were significantly abnormal in multiple rudiments at TS24, but such abnormalities partially or completely recovered in all rudiments by the prenatal stage TS27. In contrast, joint cavitation did not recover from the lack of skeletal muscle over development. Understanding how mammalian bones and joints continue to develop in an environment without muscle contractions, but with mechanical stimulation due to the movement of the mother, provides important insights into conditions affecting human babies such as developmental dysplasia of the hip and arthrogryposis. Our animal model data would suggest that the effects of immobility *in utero* may reduce in severity as development progresses.

## Acknowledgments

This work was funded by the European Research Council under the European Union’s Seventh Framework Programme (ERC Grant agreement number 336306).

